# The SARS-CoV-2 conserved macrodomain is a mono-ADP-ribosylhydrolase

**DOI:** 10.1101/2020.05.11.089375

**Authors:** Yousef M.O. Alhammad, Maithri M. Kashipathy, Anuradha Roy, Jean-Philippe Gagné, Peter McDonald, Philip Gao, Louis Nonfoux, Kevin P. Battaile, David K. Johnson, Erik D. Holmstrom, Guy G. Poirier, Scott Lovell, Anthony R. Fehr

**Author notes:** Correspondence; (K.S.).

## Abstract

Severe acute respiratory syndrome coronavirus 2 (SARS-CoV-2) and other SARS-like-CoVs encode 3 tandem macrodomains within non-structural protein 3 (nsp3). The first macrodomain, Mac1, is conserved throughout CoVs, and binds to and hydrolyzes mono-ADP-ribose (MAR) from target proteins. Mac1 likely counters host-mediated anti-viral ADP-ribosylation, a posttranslational modification that is part of the host response to viral infections. Mac1 is essential for pathogenesis in multiple animal models of CoV infection, implicating it as a virulence factor and potential therapeutic target. Here we report the crystal structure of SARS-CoV-2 Mac1 in complex with ADP-ribose. SARS-CoV-2, SARS-CoV and MERS-CoV Mac1 exhibit similar structural folds and all 3 proteins bound to ADP-ribose with low μM affinities. Importantly, using ADP-ribose detecting binding reagents in both a gel-based assay and novel ELISA assays, we demonstrated de-MARylating activity for all 3 CoV Mac1 proteins, with the SARS-CoV-2 Mac1 protein leading to a more rapid loss of substrate compared to the others. In addition, none of these enzymes could hydrolyze poly-ADP-ribose. We conclude that the SARS-CoV-2 and other CoV Mac1 proteins are MAR-hydrolases with similar functions, indicating that compounds targeting CoV Mac1 proteins may have broad anti-CoV activity.

**IMPORTANCE:** SARS-CoV-2 has recently emerged into the human population and has led to a worldwide pandemic of COVID-19 that has caused greater than 900 thousand deaths worldwide. With, no currently approved treatments, novel therapeutic strategies are desperately needed. All coronaviruses encode for a highly conserved macrodomain (Mac1) that binds to and removes ADP-ribose adducts from proteins in a dynamic post-translational process increasingly recognized as an important factor that regulates viral infection. The macrodomain is essential for CoV pathogenesis and may be a novel therapeutic target. Thus, understanding its biochemistry and enzyme activity are critical first steps for these efforts. Here we report the crystal structure of SARS-CoV-2 Mac1 in complex with ADP-ribose, and describe its ADP-ribose binding and hydrolysis activities in direct comparison to SARS-CoV and MERS-CoV Mac1 proteins. These results are an important first step for the design and testing of potential therapies targeting this unique protein domain.

## INTRODUCTION

The recently emerged pandemic outbreak of COVID-19 is caused by a novel coronavirus named severe acute respiratory syndrome coronavirus 2 (SARS-CoV-2) (1, 2). As of September 16, 2020, this virus has been responsible for ~ 30 million cases of COVID-19 and >900,000 deaths worldwide. SARS-CoV-2 is a member of the lineage B β-CoVs with overall high sequence similarity with other SARS-like CoVs, including SARS-CoV. While most of the genome is >80% similar with SARS-CoV, there are regions where amino acid conservation is significantly lower. As expected, the most divergent proteins in the SARS-CoV-2 genome from SARS-CoV include the Spike glycoprotein and several accessory proteins including 8a (absent), 8b (extended), and 3b (truncated). However, somewhat unexpectedly, several non-structural proteins also show significant divergence from SARS-CoV, including non-structural proteins 3, 4, and 7, which could affect the biology of SARS-CoV-2 (3, 4).

Coronaviruses encode 16 non-structural proteins that are translated from two open reading frames (ORFs), replicase 1a and 1ab (rep1a and rep1ab) (5). The largest non-structural protein is the non-structural protein 3 (nsp3) that encodes for multiple modular protein domains. These domains in SARS-CoV-2 diverge in amino acid sequence from SARS-CoV as much as 30%, and SARS-CoV-2 nsp3 includes a large insertion of 25-41 residues just upstream of the first of three tandem macrodomains (Mac1, Mac2, and Mac3) (Fig. 1A) (3). In addition to this insertion, the individual macrodomains show large amounts of amino acid divergence. Mac1 diverges 28% from SARS-CoV and 59% from MERS-CoV, while Mac2 and Mac3 diverge 24% from SARS-CoV. It is feasible that these significant sequence differences could impact the unique biology of SARS-CoV-2. However, macrodomains have a highly conserved structure, and thus sequence divergence may have little impact on their overall function. Mac1 is present in all CoVs, unlike Mac2 and Mac3, and early structural and biochemical data demonstrated that it contains a conserved three-layered α/β/α fold and binds to mono-ADP-ribose (MAR) and other related molecules (6–10). This is unlike Mac2 and Mac3, which fail to bind ADP-ribose and instead appear to bind to nucleic acids (11, 12). ADP-ribose is buried in a hydrophobic cleft of Mac1 where the ADP-ribose binds to several highly-conserved residues such as aspartic acid at position 23 (D23) and asparagine at position 41 (N41) of SARS-CoV (Fig. 1B) (6). Mac1 homologs are also found in alphaviruses, Hepatitis E virus, and Rubella virus, and structural analysis of these macrodomains have demonstrated that they are very similar to CoV Mac1 (13, 14). All are members of the larger MacroD-type macrodomain family, which includes human macrodomains Mdo1 and Mdo2 (15).

**Figure 1.**
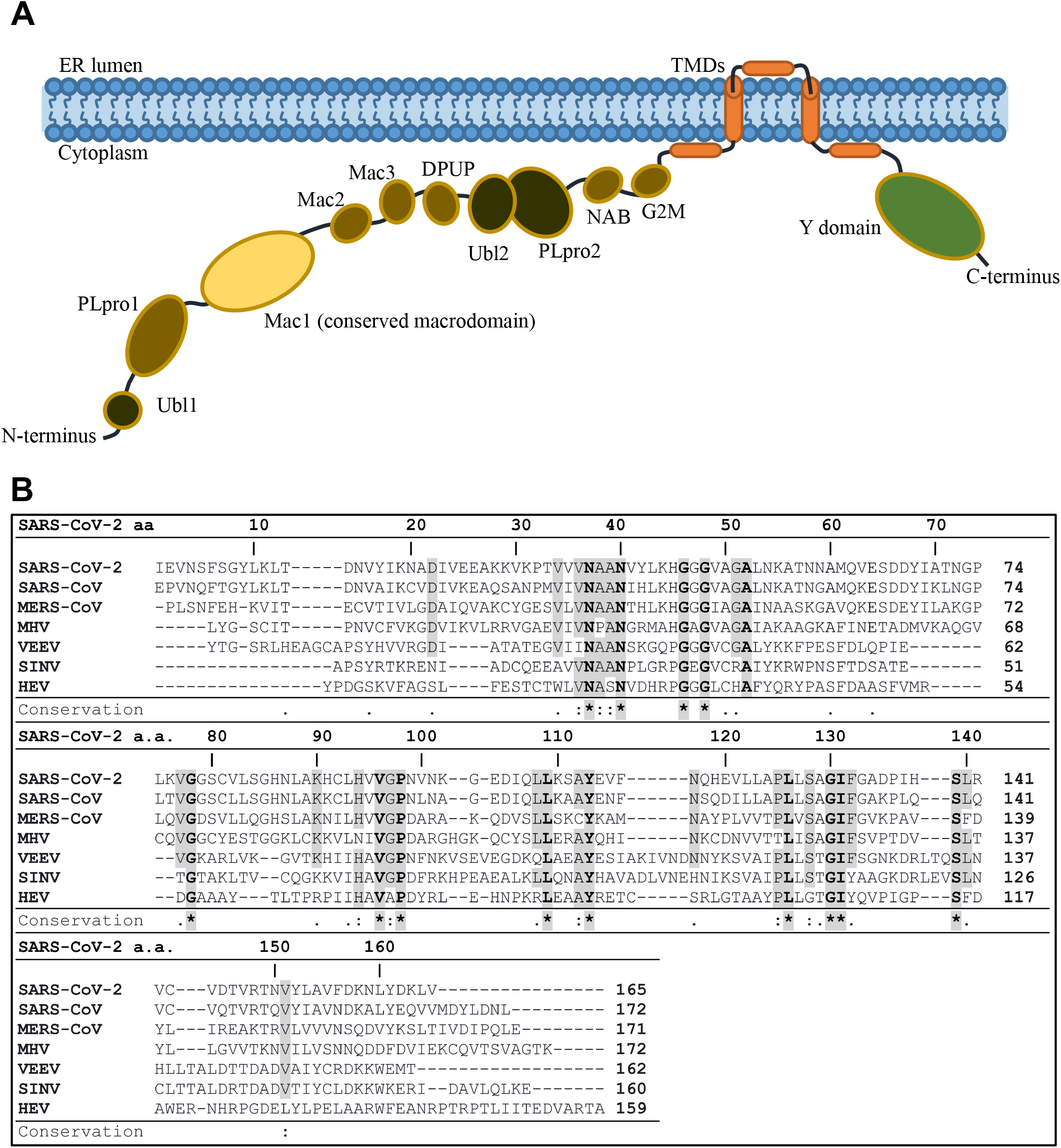
The SARS-CoV-2 Mac1 is a small domain within nsp3 and is highly conserved between other human CoV Mac1 protein domains. (A) Cartoon Schematic of the SARS-CoV-2 non-structural protein 3. The conserved macrodomain, or Mac1, is highlighted in yellow. (B) Sequence alignment of Mac1 from CoVs; SARS-CoV-2, SARS-CoV, MERS-CoV, and mouse hepatitis virus (MHV), and from alphaviruses Venezuelan equine encephalitis virus (VEEV) and sindbis virus (SINV), and hepatitis E virus (HEV). Sequences were aligned using the ClustalW method from Clustal Omega online tool with manual adjustment. Identical residues are bolded, shaded in grey, and marked with asterisks; semiconserved residues were shaded in grey and marked with two dots (one change amongst all viruses) or one dot (2 changes or conserved within CoV family).

The CoV Mac1 was originally named ADP-ribose-1”-phosphatase (ADRP) based on data demonstrating that it could remove the phosphate group from ADP-ribose-1”-phosphate (6–8). However, the activity was rather modest, and it was unclear why this would impact a virus infection. More recently it has been demonstrated that CoV Mac1 can hydrolyze the bond between amino acid chains and ADP-ribose molecules (16–18), indicating that it can reverse protein ADP-ribosylation (6, 8). ADP-ribosylation is a post-translational modification catalyzed by ADP-ribosyltransferases (ARTs, also known as PARPs) through transferring an ADP-ribose moiety from NAD^+^ onto target proteins (19). The ADP-ribose is transferred as a single unit of MAR, or single units of MAR are transferred consecutively to form a PAR chain. Several Mac1 proteins have been shown to hydrolyze MAR, but have minimal activity towards PAR (16, 17). Several MARylating PARPs are induced by interferon (IFN) and are known to inhibit virus replication, implicating MARylation in the host-response to infection (20).

Several reports have addressed the role of Mac1 on the replication and pathogenesis of CoVs, mostly using the mutation of a highly conserved asparagine to alanine (N41A-SARS-CoV). This mutation abolished the MAR-hydrolase activity of SARS-CoV Mac1 (18). This mutation has minimal effects on CoV replication in transformed cells, but reduces viral load, leads to enhanced IFN production, and strongly attenuates both murine hepatitis virus (MHV) and SARS-CoV in mouse models of infection (7, 18, 21, 22). MHV Mac1 was also required for efficient replication in primary macrophages, which could be partially rescued by the PARP inhibitors XAV-939 and 3-AB or siRNA knockdown of PARP12 or PARP14 (23). These data suggest that Mac1’s likely function is to counter PARP-mediated anti-viral ADP-ribosylation (24). Mutations in the alphavirus and HEV macrodomain also have substantial phenotypic effects on virus replication and pathogenesis (16, 25–28). As viral macrodomains are clearly important virulence factors, they are considered to be potential targets for anti-viral therapeutics (24).

Based on the close structural similarities between viral macrodomains, we hypothesized that SARS-CoV-2 Mac1 has similar binding and hydrolysis activity as other CoV Mac1 enzymes. In this study, we determined the crystal structure of the SARS-CoV-2 Mac1 protein bound to ADP-ribose. Binding to and hydrolysis of MAR was tested and directly compared to a human macrodomain (Mdo2) and the SARS-CoV and MERS-CoV Mac1 proteins by several *in vitro* assays. All CoV Mac1 proteins bound to MAR and could remove MAR from a protein substrate. However, the initial rate associated with the loss of substrate was largest for the SARS-CoV-2 Mac1 protein, especially under multi-turnover conditions. In addition, none of these enzymes could remove PAR from a protein substrate. These results indicate that Mac1 protein domains likely have similar functions, and will be instrumental in the design and testing of novel therapeutic agents targeting the CoV Mac1 protein domain.

## RESULTS

### Structure of the SARS-CoV-2 Mac1 complexed with ADP-ribose

To create recombinant SARS-CoV-2 Mac1 for structure determination and enzyme assays, nucleotides 3348-3872 of SARS-CoV-2 isolate Wuhan-hu-1 (accession number NC_045512), representing amino acids I1023-K1197 of rep1a, were cloned into a bacterial expression vector containing an N-terminal 6X-His tag and TEV cleavage site. We obtained large amounts (>100 mg) of purified recombinant protein (Fig. S1A). A small amount of this protein was digested by the TEV protease to obtain protein devoid of any extra tags for crystallization and used to obtain crystals from which the structure was determined (Fig. S1B). Our crystallization experiments resulted in the same crystal form (needle clusters) from several conditions, but only when ADP-ribose was added to the protein. This represents an additional crystal form (*P*2_1_) amongst the recently determined SARS-CoV-2 macrodomain structures (29–31).

The structure of SARS-CoV-2 Mac1 complexed with ADP-ribose was obtained using X-ray diffraction data to 2.2 Å resolution and contained four molecules in the asymmetric unit that were nearly identical. The polypeptide chains could be traced from V3-M171 for subunits A/C and V3-K172 for subunits B/D. Superposition of subunits B-D onto subunit A (169 residues aligned) yielded RMSD deviations of 0.17 Å, 0.17 Å and 0.18 Å respectively between Cα atoms. As such, subunit A was used for the majority of the structure analysis described herein. The SARS-CoV-2 Mac1 protein adopted a fold consistent with the MacroD sub-family of macrodomains that contains a core composed of a mixed arrangement of 7 β-sheets (parallel and antiparallel) that are flanked by 6 α-helices (Fig. 2A-B).

**Figure 2.**
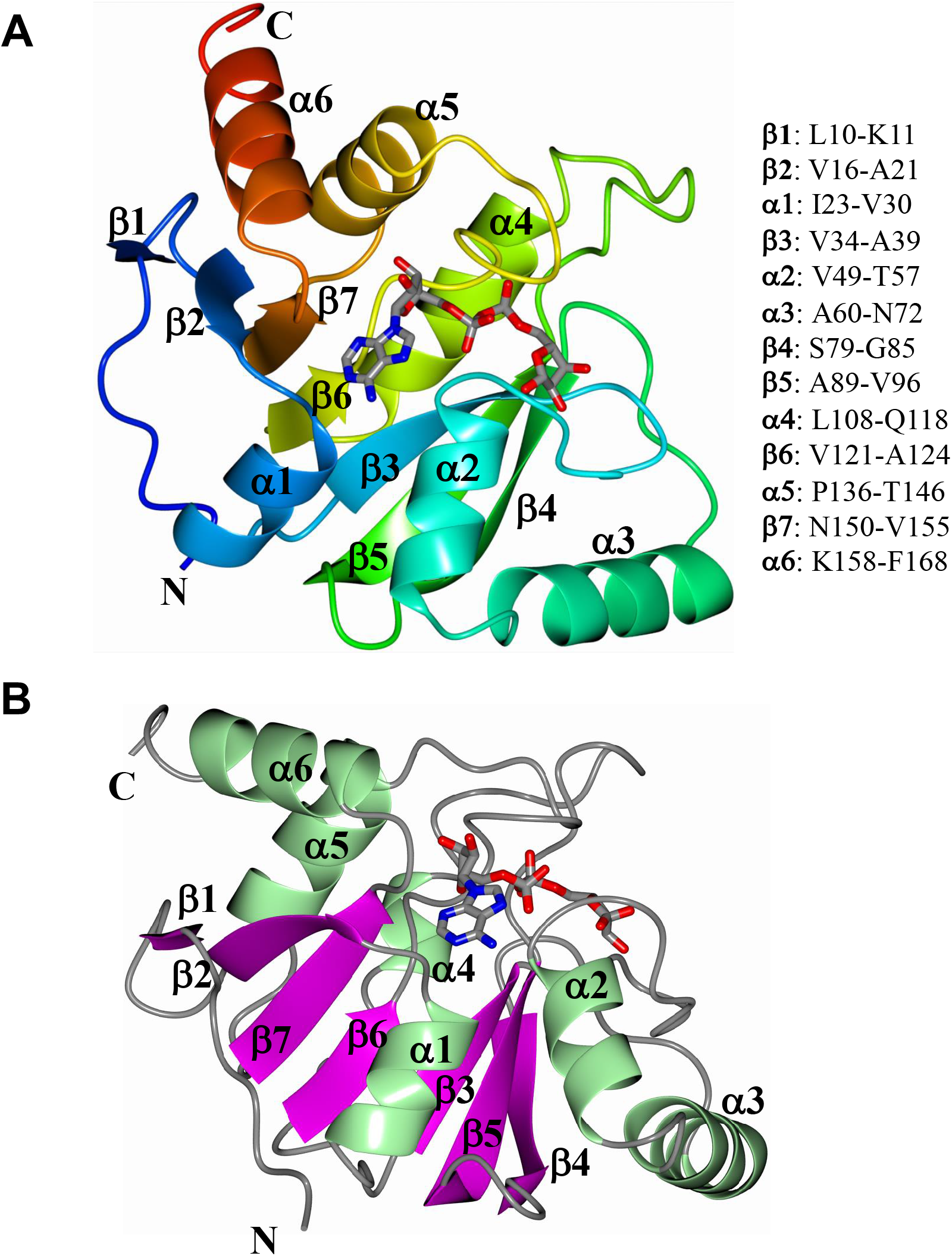
Structure of SARS-CoV-2 Mac1 complexed with ADP-ribose. **A)** The structure was rendered as a blend through model from the N-terminus (blue) to the C-terminus (red). **B)** The structure was colored by secondary structure showing sheets (magenta) and helices (green). The ADP-ribose is rendered as gray cylinders with oxygens and nitrogens colored red and blue, respectively.

As mentioned above, apo crystals were never observed for our construct, though the apo structure has been solved by researchers at The Center for Structural Genomics of Infectious Diseases (PDB 6WEN) (30) and the University of Wisconsin-Milwaukee (PDB 6WEY) (32). Further analysis of the amino acid sequences used for expression and purification revealed that our construct had 5 additional residues at the C-terminus (MKSEK) and differs slightly at the N-terminus as well (GIE vs GE) relative to 6WEN. In addition, the sequence used to obtain the structure of 6WEY is slightly shorter than SARS-CoV-2 Mac1 at both the N and C-terminal regions (Fig. S2A). To assess the effect of these additional residues on crystallization, chain B of the SARS-CoV-2 Mac1, which was traced to residue K172, was superimposed onto subunit A of PDB 6W02 (31), a previously determined structure of ADP-ribose bound SARS-CoV-2 Mac1. Analysis of the crystal packing of 6W02 indicates that the additional residues at the C-terminus would clash with symmetry related molecules (Fig. S2B). This suggests that the presence of these extra residues at the C-terminus likely prevented the generation of the more tightly packed crystal forms obtained for 6W02 and 6WEY, which diffracted to high resolution.

The ADP-ribose binding pocket contained large regions of positive electron density consistent with the docking of ADP-ribose (Fig. 3A). The adenine forms two hydrogen bonds with D22-I23, which makes up a small loop between β2 and the N-terminal half of α1. The side chain of D22 interacts with N6, while the backbone nitrogen atom of I23 interacts with N1, in a very similar fashion to the SARS-CoV macrodomain (6). This aspartic acid is known to be critical for ADP-ribose binding for alphavirus macrodomains (26, 27). A large number of contacts are made in the highly conserved loop between β3 and α2 which includes many highly-conserved residues, including a GGG (motif) and N40, which is completely conserved in all enzymatically active macrodomains (33). N40 is positioned to make hydrogen bonds with the 3’ OH groups of the distal ribose, as well as a conserved water molecule (Fig. 3B-C). K44 and G46 also make hydrogen bonds with the 2’ OH of the distal ribose, G48 makes contact with the 1’ OH and a water that resides near the catalytic site, while the backbone nitrogen atom of V49 hydrogen bonds with the α-phosphate. The other major interactions with ADP-ribose occur in another highly conserved region consisting of residues G130, I131, and F132 that are in the loop between β6 and α5 (Fig. 3B). The α-phosphate accepts a hydrogen bond from the nitrogen atom of I131, while the β-phosphate accepts hydrogen bonds from the backbone nitrogen atom of G130 and F132. The phenyl ring of F132 may make van der Waals interactions with the distal ribose to stabilize it, which may contribute to binding and hydrolysis (34). Loops β3-α2 and β6-α5 are connected by an isoleucine bridge that forms a narrow channel around the diphosphate which helps position the terminal ribose for water-mediated catalysis (6). Of all these residues, is not exactly clear which ones are important for ADP-ribose binding, hydrolysis, or both. Additionally, a network of direct contacts of ADP-ribose to solvent along with water mediated contacts to the protein are shown (Fig. 3C).

**Figure 3.**
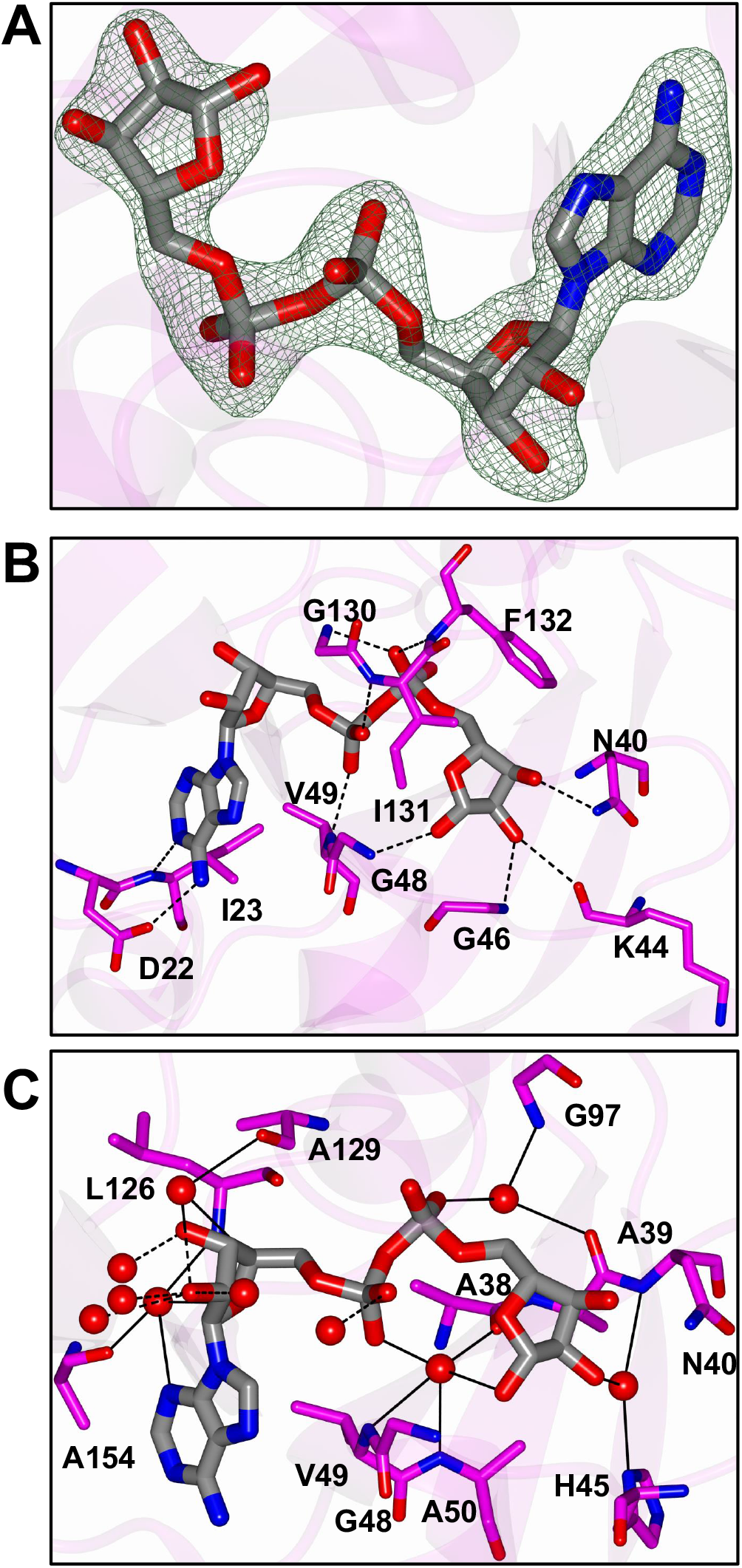
Binding mode of ADP-ribose in SARS-CoV-2 Mac1. **A)** Fo-Fc Polder omit map (green mesh) contoured at 3σ. **B)** Hydrogen bond interactions (dashed lines) between ADP-ribose and amino acids. **C)** Interactions with water molecules. Direct hydrogen bond interactions are represented by dashed lines and water mediated contacts to amino acids are drawn as solid lines.

### Comparison of SARS-CoV-2 Mac1 with other CoV macrodomain structures

We next sought to compare the SARS-CoV-2 Mac1 to other deposited structures of this protein. Superposition with Apo (6WEN) and ADP-ribose complexed protein (6W02) yielded RMSD of 0.48 Å (168 residues) and 0.37 Å (165 residues), respectively, indicating a high degree of similarity (Fig. S3A-B). Comparison of the ADP-ribose binding site of SARS-CoV-2 Mac1 with that of the apo structure (6WEN) revealed minor conformational differences in order to accommodate ADP-ribose binding. The loop between β3 and α2 (H45-V49) undergoes a change in conformation and the sidechain of F132 is moved out of the ADP-ribose binding site (Fig. S3C). Our ADP-ribose bound structure is nearly identical to 6W02, except for slight deviations in the β3-α2 loop and an altered conformation of F156, where the aryl ring of F156 is moved closer to the adenine ring (Fig. S3 C-D). However, this is likely a result of crystal packing as F156 adopts this conformation in each subunit and would likely clash with subunit residues related by either crystallographic or non-crystallographic symmetry.

We next compared the ADP-ribose bound SARS-CoV-2 Mac1 structure with that of SARS-CoV (PDB 2FAV) (6) and MERS-CoV (PDB 5HOL) (35) Mac1 proteins. Superposition yielded RMSD deviations of 0.71 Å (166 residues) and 1.06 Å (161 residues) for 2FAV and 5HOL, respectively. Additionally, the ADP-ribose binding mode in the SARS-CoV and SARS-CoV-2 structures almost perfectly superimposed (Fig. 4A-D). The conserved aspartic acid residue (D22, SARS-CoV-2) that binds to adenine, is localized in a similar region in all 3 proteins although there are slight differences in the rotamers about the Cβ-Cγ bond. The angles between the mean planes defined by the OD1, CG and OD2 atoms relative to SARS-CoV-2 Mac1 is 23.1° and 46.5° for the SARS-CoV and MERS-CoV Mac1 structures, respectively. Another notable difference is that SARS-CoV and SARS-CoV-2 macrodomains have an isoleucine (I23) following this aspartic acid while MERS-CoV has an alanine (A22). Conversely, SARS-CoV-2 and SARS-CoV Mac1 have a valine instead of an isoleucine immediately following the GGG motif (V49/I48). From these structures it appears that having two isoleucines in this location would clash, and that lineage B and lineage C β-CoVs has evolved in unique ways to create space in this pocket (Fig. 4D and data not shown). Despite these small differences in local structure, the overall structure of CoV Mac1 domains remain remarkably conserved, and indicates they likely have similar biochemical activities and biological functions.

**Figure 4.**
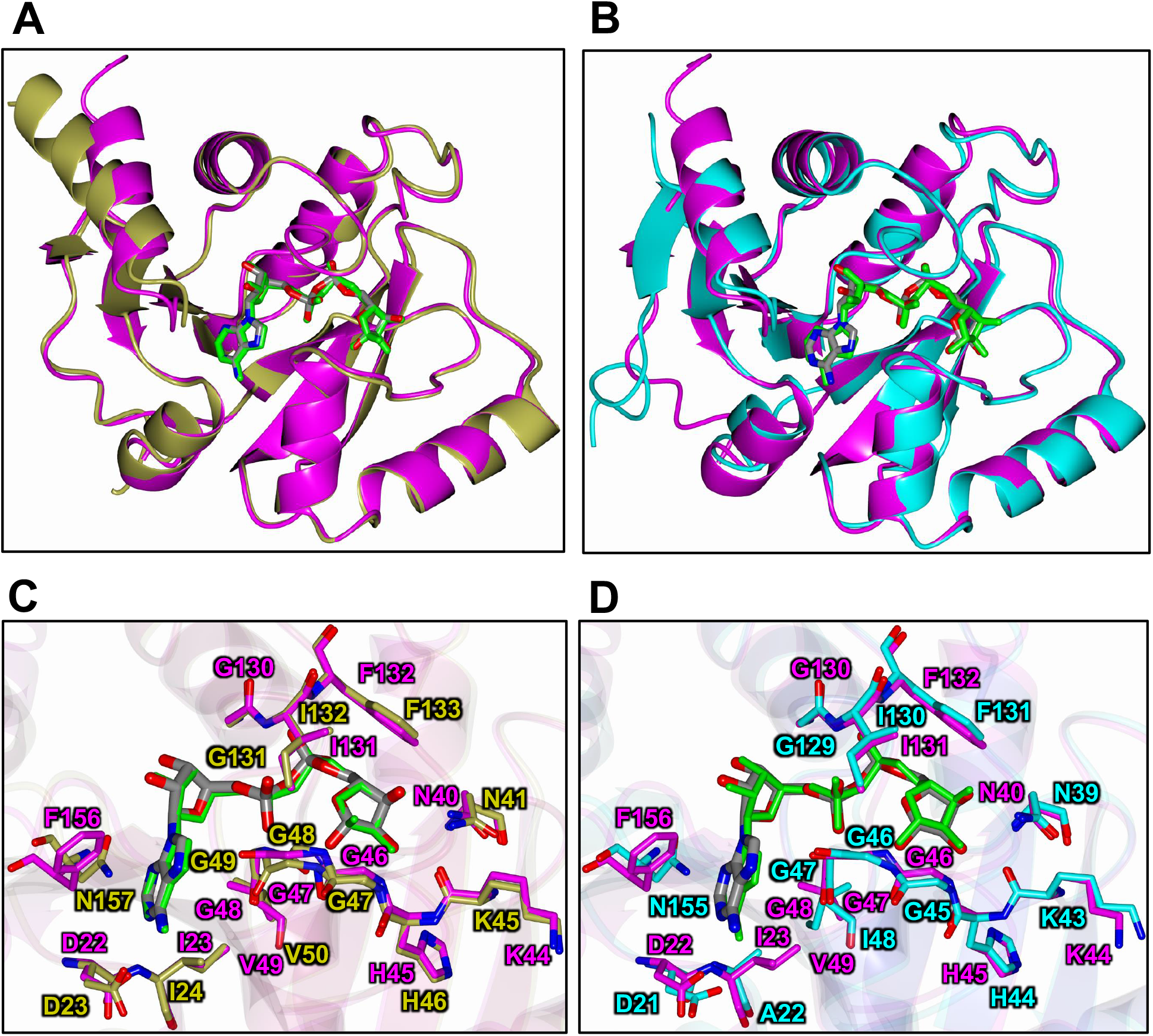
Structural comparison of the SARS-CoV-2 Mac1 protein with the SARS-CoV and MERS-CoV Mac1 proteins. **A-B)** Superposition of SARS-CoV-2 macrodomain (magenta) with coronavirus macrodomain structures. **A**) SARS-CoV Mac1 with ADP-ribose (gold) (2FAV) and **B**) MERS-CoV Mac1 with ADP-ribose (teal) (5HOL). **C-D)** Superposition of SARS-CoV-2 Mac1 (magenta) with other coronavirus Mac1 structures highlighting the ADP-ribose binding site. **C**) SARS-CoV (gold), **D**) MERS-CoV (teal). The ADP-ribose molecules are colored gray for SARS-CoV-2 Mac1 (**A-D**) and are rendered as green cylinders for SARS-CoV Mac1 (panel **A,C**) and MERS-CoV Mac1 (panel **B,D**).

### SARS-CoV, SARS-CoV-2, and MERS-CoV bind to ADP-ribose with similar affinities

To determine if the CoV macrodomains had any noticeable differences in their ability to bind ADP-ribose, we performed isothermal titration calorimetry (ITC), which measures the energy released or absorbed during a binding reaction. Macrodomain proteins from human (Mdo2), SARS-CoV, MERS-CoV, and SARS-CoV-2 were purified (Fig. S1A) and tested for their affinity to ADP-ribose. All CoV Mac1 proteins bound to ADP-ribose with low micromolar affinity (7-16 μM), while human Mdo2 bound with an affinity about 10-times stronger (~220 nM) (Fig. 5A-B). As a control we tested the ability of the MERS-CoV macrodomain to bind to ATP, and only observed minimal binding with mM affinity (data not shown). At higher concentrations, the SARS-CoV-2 macrodomain caused a slightly endothermic reaction, potentially the result of protein aggregation or a change in conformation (Fig. 5A). The MERS-CoV Mac1 had a greater affinity for ADP-ribose than SARS-CoV or SARS-CoV-2 Mac1 in the ITC assay (Fig. 5A-B), however, our results found the differences between these macrodomain proteins to be much closer than previously reported (9). As an alternate method to confirm ADP-ribose binding, we conducted a thermal shift assay. All 4 macrodomains tested denatured at higher temperatures with the addition of ADP-ribose (Fig. S4). We conclude that lineage B and lineage C β-CoV Mac1 proteins bind to ADP-ribose with similar affinities.

**Figure 5.**
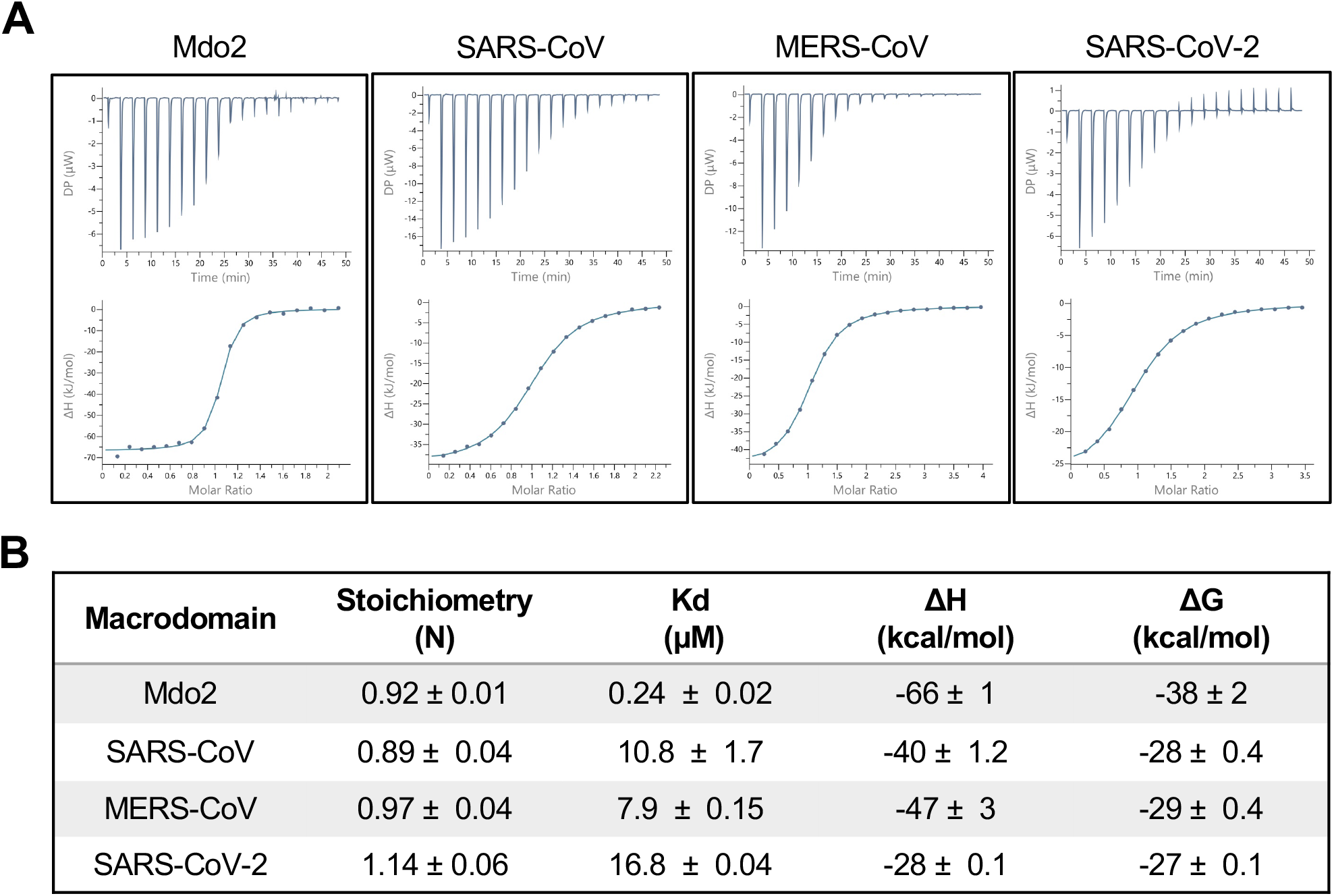
Human CoVs bind to ADP-ribose with similar affinity. **A-B)** ADP-ribose binding of human Mdo2 and SARS-CoV, MERS-CoV, and SARS-CoV-2 Mac1 proteins by ITC. Images in **(A)** are of one experiment representative of at least 2 independent experiments. Data in **(B)** represent the combined averages of multiple independent experiments for each protein. Mdo2 n=2; SARS-CoV n=5; MERS-CoV n=6; SARS-CoV-2 n=2.

### CoV macrodomains are MAR-hydrolases

To examine the MAR-hydrolase activity of CoV Mac1, we first tested the viability of using ADP-ribose binding reagents to detect MARylated protein. Previously, radiolabeled NAD^+^ has been the primary method used to label MARylated protein (16, 17). To create a MARylated substrate, the catalytic domain of the PARP10 (GST-PARP10 CD) protein was incubated with NAD^+^, leading to its automodification. PARP10 CD is a standard substrate that has been used extensively in the field to analyze the activity of macrodomains (16, 18, 26, 27). PARP10 is highly upregulated upon CoV infection (23, 36) and is known to primarily auto-MARylate itself on acidic residues, which are the targets of the MacroD2 class of macrodomains (27). We then tested a panel of anti-MAR, anti-PAR, or both anti-MAR and anti-PAR binding reagents/antibodies for the ability to detect MARylated PARP10 by immunoblot. The anti-MAR and anti-MAR/PAR binding reagents, but not anti-PAR antibody, bound to MARylated PARP10 (Fig. S5). Therefore, in this work we utilized the anti-MAR binding reagent to detect MARylated PARP10.

We next tested the ability of SARS-CoV-2 Mac1 to remove ADP-ribose from MARylated PARP10. SARS-CoV-2 Mac1 and MARylated PARP10 were incubated at equimolar amounts of protein at 37°C and the reaction was stopped at 5, 10, 20, 30, 45 or 60 minutes (Fig. 6A). As a control, MARylated PARP10 was incubated alone at 37°C and collected at similar time points (Fig. 6A and Fig. S6). Each reaction had equivalent amounts of MARylated PARP10 and Mac1 which was confirmed by Coomassie Blue staining (Fig. 6A). An immediate reduction of more than 50% band intensity was observed within five minutes, and the ADP-ribose modification was nearly completely removed by SARS-CoV-2 Mac1 within 30 minutes (Fig. 6A). The MARylated PAPR10 bands intensities were calculated, plotted, and were fit using non-linear regression (Fig. 6B). This result indicates that the SARS-CoV-2 Mac1 protein is a mono-ADP-ribosylhydrolase enzyme.

**Figure 6.**
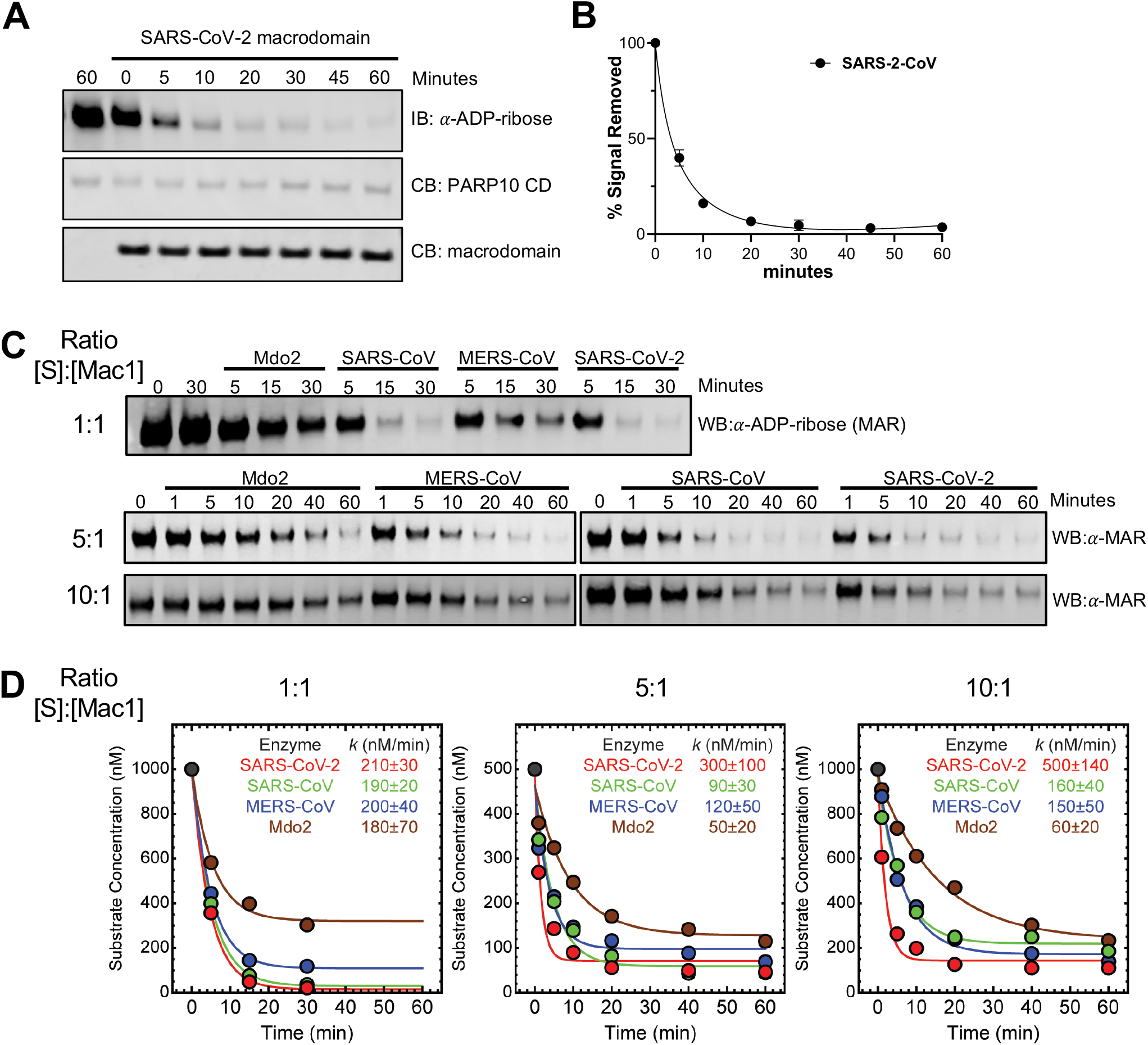
Coronavirus Mac1 proteins are ADP-ribosylhydrolases. **A)** The SARS-CoV-2 macrodomain was incubated with MARylated PARP10 CD *in vitro* at equimolar ratios (1 μM) for the indicated times at 37°C. ADP-ribosylated PARP10 CD was detected by immunoblot (IB) with anti-ADP-ribose binding reagent (Millipore-Sigma MAB1076). Total PARP10 CD and macrodomain protein levels were determined by Coomassie Blue (CB) staining. PARP10 CD incubated alone at 37°C was stopped at 0 or 60 minutes. **B)** The level of de-MARylation was measured by quantifying band intensity using Image J software. Intensity values were plotted and fit to a non-linear regression curve with error bars representing standard deviation. Results in **A** are representative experiments of two independent experiments and data in **B** represent the combined results of the two independent experiments. **C)** The Mdo2, MERS-CoV, SARS-CoV, and SARS-CoV-2 macrodomains were incubated with MARylated PARP10 CD *in vitro* at the following ratios of [substrate]:[Mac1]: 1:1 (1 μM), 5:1 (500 nM, 100 nM), or 10:1 (1 μM, 100 nM) for the indicated times at 37°C. ADP-ribosylated PARP10 CD was detected as described above, and total PARP10 CD and macrodomain protein levels were determine by Coomassie Blue (Fig. S6). **D)** Time-dependent substrate concentrations were determined by quantifying band intensity using Image Studio software. The data were then analyzed using Mathematica 12, as described in Methods, to determine the initial rate (*k*) of substrate decay. Results in **C** are representative experiments of three independent experiments and data in **D** represent the combined results of the three independent experiments.

Next, we compared the MAR-hydrolase activity of Mac1 proteins from SARS-CoV-2, SARS-CoV, and MERS-CoV and human (i.e., Mdo2). Specifically, we monitored the time-dependent loss of substrate using immunoblotting (Fig. 6C) under equimolar (i.e., 1 μM [Mac1]:1 μM [substrate]) and multiple-turnover conditions (i.e., 0.5 μM [substrate]:0.1 μM [Mac1] and 1.0 μM [substrate]:0.1 μM [Mac1]), with total protein amounts confirmed by Coomassie blue staining (Fig. S7). The resulting substrate decay plots (Fig. 6D) were fit using non-linear regression to determine the initial rate (*k*) of substrate decay. Our results indicate that the three CoV Mac1 proteins give rise to similar, but not identical, values of *k* (Fig. 6D). The SARS-CoV-2 Mac1 protein has a greater *k* than the SARS-CoV or MERS-CoV Mac1 proteins, especially under multiple-turnover conditions, and all 3 viral macrodomains gave rise to a more rapid loss of substrate than the human Mdo2 enzyme (Fig. 6F). However, further enzymatic analyses of these proteins are warranted to more thoroughly understand their kinetics and binding affinities associated with various MARylated substrates.

### CoV Mac1 proteins do not hydrolyze PAR

To determine if the CoV Mac1 proteins could remove PAR from proteins, we incubated these proteins with an auto-PARylated PARP1 protein. PARP1 was incubated with increasing concentrations of NAD^+^ to create a range of modification levels (Fig. S8). We incubated both partially and heavily modified PARP1 with all four macrodomains and PARG as a positive control for 1 hour. While PARG completely removed PAR, none of the macrodomain proteins removed PAR chains from PARP1 (Fig. 7). We conclude that macrodomain proteins are unable to remove PAR from an automodified PARP1 protein under these conditions.

**Fig. 7.**
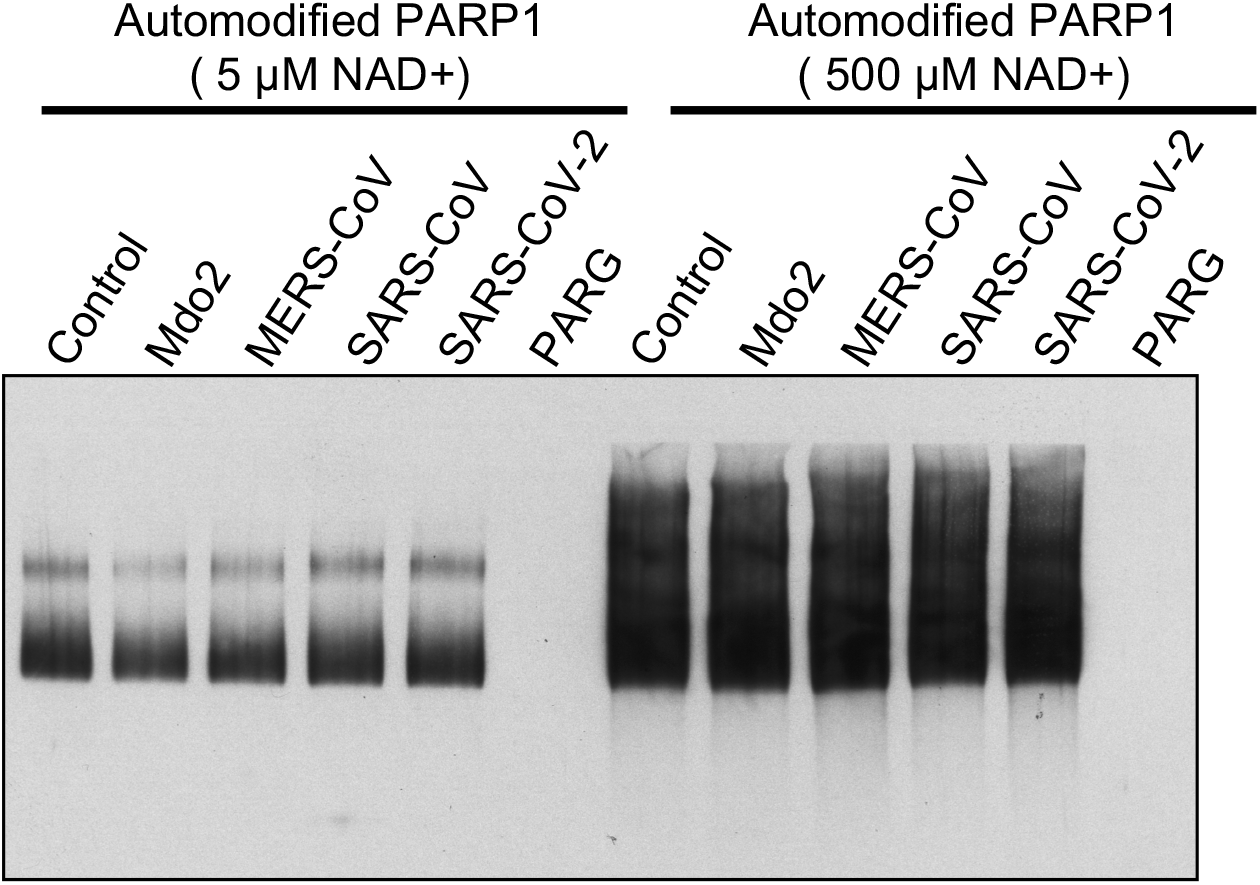
Coronavirus Mac1 proteins do not hydrolyze PAR. PAR hydrolase assays were performed with PARP1 either extensively poly-ADP-ribosylated (500 μM NAD^+^) or partially poly-ADP-ribosylated (5 μM NAD^+^) to produce oligo-ADP-ribose. Macrodomains were incubated with both automodified PARP1 substrates for 1 hour. PAR was detected by immunoblot with the anti-PAR antibody 96-10. PARG (catalytically active 60 kD fragment) was used as a positive control. The results are representative of 2 independent experiments.

### ELISA assays can be used to measure ADP-ribosylhydrolase activity of macrodomains

Gel based assays as described above suffer from significant limitations in the number of samples that can be done at once. A higher throughput assay will be needed to more thoroughly investigate the activity of these enzymes and to screen for inhibitor compounds. Based on the success of our antibody-based detection of MAR, we developed an ELISA assay that has a similar ability to detect de-MARylation as our gel-based assay, but with the ability to do so in a higher throughput manner (Fig. 8A). First, MARylated PARP10 was added to ELISA plates. Next, the wells were washed and then incubated with different concentrations of the SARS-CoV-2 Mac1 protein for 60 min. After incubation, the wells were washed and treated with anti-MAR binding reagent, followed by HRP-conjugated secondary antibody and the detection reagent. As controls, we detected MARylated and non-MARylated PARP10 proteins bound to glutathione plates with anti-GST antibody and anti-MAR binding reagents and their corresponding secondary antibodies (Fig. 8B). SARS-CoV-2 Mac1 was able to remove MAR signal in a dose-dependent manner and plotted to a linear non-regression fitted line (Fig. 8C). Based on these results, we believe that this ELISA assay will be a useful tool for screening potential inhibitors of macrodomain proteins.

**Figure 8.**
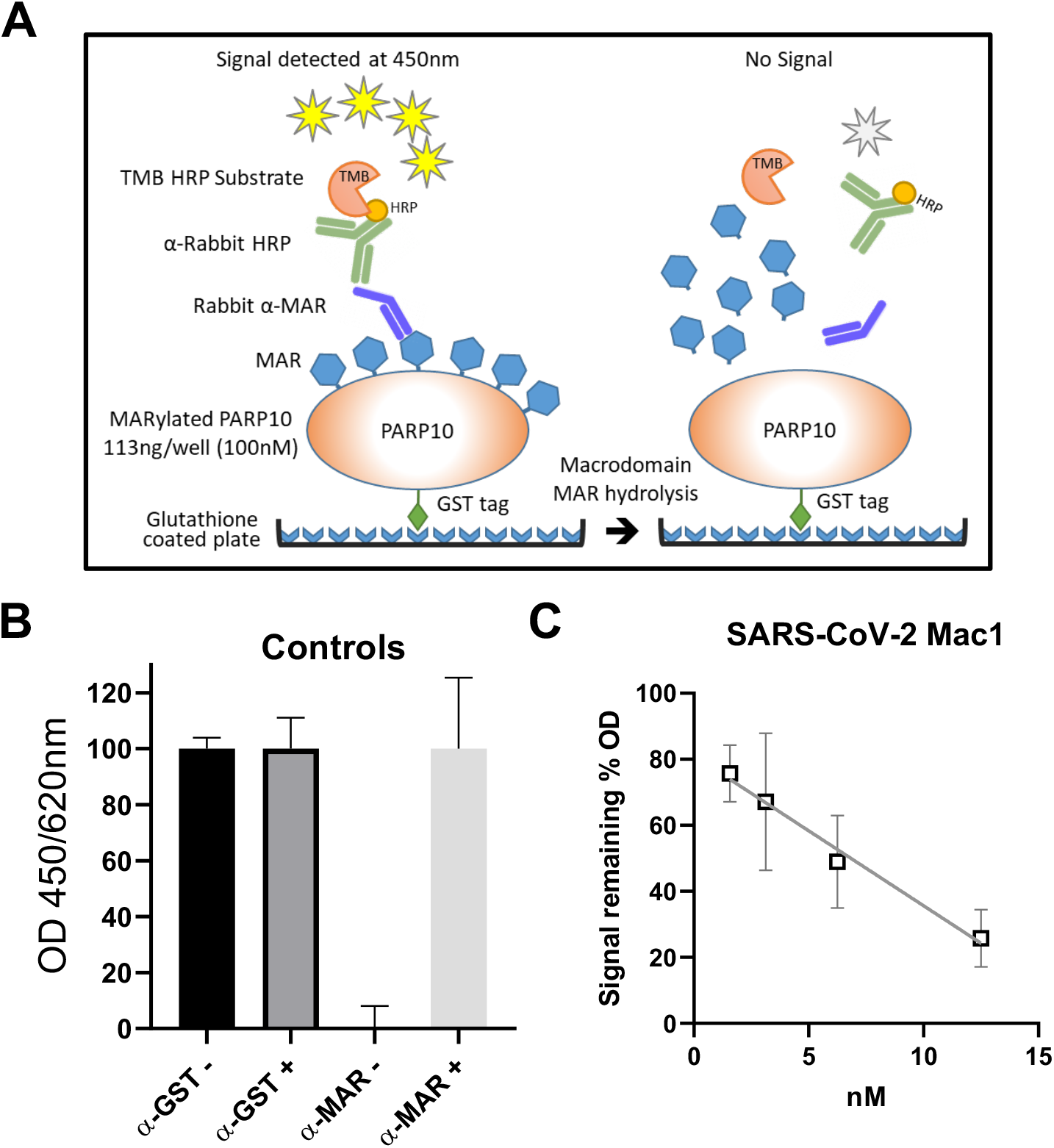
Development of an ELISA assay to detect de-MARylation. **A)** Cartoon schematic of the ELISA assay. ELISA plates pre-coated with glutathione and pre-blocked were used capture GST-tagged PARP10 proteins, which was used as a substrate for de-MARylation. The removal of MAR was detected by anti-MAR antibodies. **B)** MARylated PARP10 (MAR+) and non-MARylated PARP10 (MAR-) with no SARS-CoV-2 Mac1 as controls were detected with anti-mono-ADP-ribose binding reagent (α-MAR) (Millipore-Sigma MAB1076) or with anti-GST (α-GST) (Invitrogen, MA4-004). **C)** Starting at 12.5 nM, 2-fold serial dilutions of the SARS-CoV-2 Mac1 protein was incubated in individual wells with MARylated PARP10-CD for 60 min at 37°C. The graph represents the combined results of 2 independent experiments.

## DISCUSSION

Here we report the crystal structure of SARS-CoV-2 Mac1 and its enzyme activity *in vitro*. Structurally, it has a conserved three-layered α/β/α fold typical of the MacroD family of macrodomains, and is extremely similar to other CoV Mac1 proteins (Fig. 2–4). The conserved CoV macrodomain (Mac1) was initially described as an ADP-ribose-1”-phosphatase (ADRP), as it was shown to be structurally similar to yeast enzymes that have this enzymatic activity (37). Early biochemical studies confirmed this activity for CoV Mac1, though its phosphatase activity for ADP-ribose-1”-phosphate was rather modest (6–8). Later, it was shown that mammalian macrodomain proteins could remove ADP-ribose from protein substrates, indicating protein de-ADP-ribosylation as a more likely function for the viral macrodomains (33, 38, 39). Shortly thereafter, the SARS-CoV, hCoV-229E, FIPV, several alphavirus, and the hepatitis E virus macrodomains were demonstrated to have de-ADP-ribosylating activity (16–18). However, this activity has not yet been reported for the MERS-CoV or SARS-CoV-2 Mac1 protein.

In this study, we show that the Mac1 proteins from SARS-CoV, MERS-CoV and SARS-CoV-2 hydrolyze MAR from a protein substrate (Fig. 6). Their enzymatic activities were similar despite sequence divergence of almost 60% between SARS-CoV-2 and MERS-CoV. However, the initial rate associated with the loss of substrate was largest for the SARS-CoV-2 Mac1 protein, particularly under multiple-turnover conditions. It is unclear what structural or sequence differences may account for the increased activity of the SARS-CoV-2 Mac1 protein under these conditions, especially considering the pronounced structurally similarities between these proteins, specifically the SARS-CoV Mac1 (0.71 Å RMSD). It is also unclear if these differences would matter in the context of the virus infection, as the relative concentrations of Mac1 and its substrate during infection is not known. We also compared these activities to the human Mdo2 macrodomain. Mdo2 had a greater affinity for ADP-ribose than the viral enzymes, but had significantly reduced enzyme activity in our experiments. Due to its high affinity for ADP-ribose, it is possible that the Mdo2 protein was partially inhibited by rebinding to the MAR product in these assays. Regardless, these results suggest that the human and viral proteins likely have structural differences that alter their biochemical activities *in vitro*, indicating that it may be possible to create viral macrodomain inhibitors that don’t impact the human macrodomains. We also compared the ability of these macrodomain proteins to hydrolyze PAR. None of the macrodomains were able to hydrolyze either partially or heavily modified PARP1, further demonstrating that the primary enzymatic activity of these proteins is to hydrolyze MAR (Fig. 7).

When analyzing viral macrodomain sequences, it is clear that they have at least 3 highly conserved regions (Fig. 1B)(24). The first region includes the NAAN (37–40) and GGG (residues 46-48) motifs in the loop between β3 and α2. The second domain includes a GIF (residues 130-132) motif in the loop between β6 and α5. The final conserved region is a VGP (residues 96-98) motif at the end of β5 and extends into the loop between β5 and α4. Both of the first two domains have well defined interactions with ADP-ribose (Fig. 3). However, no one has addressed the role of the VGP residues, though our structure indicates that the glycine may interact with a water molecule that makes contact with the β-phosphate. Identifying residues that directly contribute to ADP-ribose binding, hydrolysis, or both by CoV Mac1 proteins will be critical to determining the specific roles of ADP-ribose binding and hydrolysis in CoV replication and pathogenesis.

While all previous studies of macrodomain de-ADP-ribosylation have primarily used radiolabeled substrate, we obtained highly repeatable and robust data utilizing ADP-ribose binding reagents designed to specifically recognize MAR (40, 41). The use of these binding reagents should enhance the feasibility of this assay for many labs that are not equipped for radioactive work. Utilizing these binding reagents, we further developed an ELISA assay for de-MARylation that has the ability to dramatically increase the number of samples that can be analyzed compared to the gel-based assay. To our knowledge, previously developed ELISA assays were used to measure ADP-ribosyltransferase activities (42) but no ELISA has been established to test the ADP-ribosylhydrolase activity of macrodomain proteins. This ELISA assay should be useful to those in the field to screen compounds for macrodomain inhibitors that could be either valuable research tools or potential therapeutics.

The functional importance of the CoV Mac1 domain has been demonstrated in several reports, mostly utilizing the mutation of a highly conserved asparagine that mediates contact with the distal ribose (Fig. 3B) (18, 21, 22). However, the physiological relevance of Mac1 during SARS-CoV-2 infection has yet to be determined. In addition, the proteins that are targeted by the CoV Mac1 for de-ADP-ribosylation remains unknown. Unfortunately, there are no known compounds that inhibit this domain that could help identify the functions of this protein during infection. The outbreak of COVID-19 has illustrated an urgent need for developing multiple therapeutic drugs targeting conserved coronavirus proteins. Mac1 appears to be an ideal candidate for further drug development based on: *i)* its highly conserved structure and biochemical activities within CoVs; and *ii)* its importance for multiple CoVs to cause disease.

Targeting Mac1 may also have the benefit of enhancing the innate immune response, as we have shown that Mac1 is required for some CoVs to block IFN production (18, 23). Considering that Mac1 proteins from divergent αCoVs such as 229E and FIPV also have de-ADP-ribosylating activity (16, 17), it is possible that compounds targeting Mac1 could prevent disease caused by of wide variety of CoV, including those of veterinary importance like porcine epidemic diarrhea virus (PEDV). Additionally, compounds that inhibit Mac1 in combination with the structure could help identify the mechanisms it uses to bind to its biologically relevant protein substrates, remove ADP-ribose from these proteins, and potentially define the precise function for Mac1 in SARS-CoV-2 replication and pathogenesis. In conclusion, the results described here will be critical for the design and development of highly-specific Mac1 inhibitors that could be used therapeutically to mitigate COVID-19 or future CoV outbreaks.

## METHODS

### Plasmids

The SARS-CoV macrodomain (Mac1) (residues 1000-1172 of pp1a) was cloned into the pET21a+ expression vector with an N-terminal His tag. The MERS-CoV Mac1 (residues 1110-1273 of pp1a) was also cloned into pET21a+ with a C-terminal His tag. SARS-CoV-2 Mac1 (residues 1023-1197 of pp1a) was cloned into the pET30a+ expression vector with an N-terminal His tag and a TEV cleavage site (Synbio). The pETM-CN Mdo2 Mac1 (residues 7-243) expression vector with an N-terminal His-TEV-V5 tag and the pGEX4T-PARP10-CD (residues 818-1025) expression vector with an N-terminal GST tag were previously described (33). All plasmids were confirmed by restriction digest, PCR, and direct sequencing.

### Protein Expression and Purification

A single colony of *E. coli* cells (C41(DE3)) containing plasmids harboring the constructs of the macrodomain proteins was inoculated into 10 mL LB media and grown overnight at 37°C with shaking at 250 rpm. The overnight culture was transferred to a shaker flask containing 2X 1L TB media at 37°C until the OD600 reached 0.7. The proteins were either induced with 0.4 mM IPTG at 37°C for 3 hours, or 17°C for 20 hours. Cells were pelleted at 3500 × g for 10 min and frozen at −80°C. Frozen cells were thawed at room temperature, resuspended in 50 mM Tris (pH 7.6), 150 mM NaCl, and sonicated using the following cycle parameters: Amplitude: 50%, Pulse length: 30 seconds, Number of pulses: 12, while incubating on ice for >1min between pulses. The soluble fraction was obtained by centrifuging the cell lysate at 45,450 × g for 30 minutes at 4°C. The expressed soluble proteins were purified by affinity chromatography using a 5 ml prepacked HisTrap HP column on an AKTA Pure protein purification system (GE Healthcare). The fractions were further purified by size-exclusion chromatography (SEC) with a Superdex 75 10/300 GL column equilibrated with 20mM Tris (pH 8.0), 150 mM NaCl and the protein sized as a monomer relative to the column calibration standards. To cleave off the His tag from the SARS-CoV-2 Mac1, purified TEV protease was added to purified SARS-CoV-2 Mac1 protein at a ratio of 1:10 (w/w), and then passed back through the Ni-NTA HP column. Protein was collected in the flow through and equilibrated with 20 mM Tris (pH 8.0), 150 mM NaCl. The SARS-CoV-2 Mac1, free from the N-terminal 6X-His tag, was used for subsequent crystallization experiments.

For the PARP10-CD protein, the cell pellet was resuspended in 50 mM Tris-HCl (pH 8.0), 500 mM NaCl, 0.1mM EDTA, 25% glycerol, 1 mM DTT and sonicated as described above. The cell lysate was incubated with 10 ml of Glutathione Sepharose 4B resin from GE Healthcare, equilibrated with the same buffer for 2 hours, then applied to a gravity flow column to allow unbound proteins to flow through. The column was washed with the resuspension buffer till the absorbance at 280 nm reached baseline. The bound protein was eluted out of the column with resuspension buffer containing 20 mM reduced glutathione and then dialyzed back into the resuspension buffer overnight at 4°C.

### Isothermal Titration Calorimetry

All ITC titrations were performed on a MicroCal PEAQ-ITC instrument (Malvern Pananalytical Inc., MA). All reactions were performed in 20 mM Tris pH 7.5, 150 mM NaCl using 100 μM of all macrodomain proteins at 25°C. Titration of 2 mM ADP-ribose or ATP (MilliporeSigma) contained in the stirring syringe included a single 0.4 μL injection, followed by 18 consecutive injections of 2 μL. Data analysis of thermograms was analyzed using one set of binding sites model of the MicroCal ITC software to obtain all fitting model parameters for the experiments.

### Differential Scanning Fluorimetry (DSF)

Thermal shift assay with DSF involved use of LightCycler® 480 Instrument (Roche Diagnostics). In total, a 15 μL mixture containing 8X SYPRO Orange (Invitrogen), and 10 μM macrodomain protein in buffer containing 20 mM Hepes, NaOH, pH 7.5 and various concentrations of ADP-ribose were mixed on ice in 384-well PCR plate (Roche). Fluorescent signals were measured from 25 to 95 °C in 0.2 °C/30-s steps (excitation, 470-505 nm; detection, 540-700 nm). The main measurements were carried out in triplicate. Data evaluation and Tm determination involved use of the Roche LightCycler® 480 Protein Melting Analysis software, and data fitting calculations involved the use of single site binding curve analysis on Graphpad Prism.

### MAR Hydrolase Assays

#### Automodification of PARP10-CD protein

A 10 μM solution of purified PAPR10-CD protein was incubated for 20 minutes at 37°C with 1 mM final concentration of β-Nicotinamide Adenine Dinucleotide (β NAD^+^) (Millipore-Sigma) in a reaction buffer (50 mM HEPES, 150 mM NaCl, 0.2 mM DTT, and 0.02% NP-40). MARylated PARP10 was aliquoted and stored at − 80°C.

#### PAPR10-CD ADP-ribose hydrolysis

All reactions were performed at 37°C for the designated time. A 1 μM solution of MARylated PARP10-CD and purified Mac1 protein was added in the reaction buffer (50 mM HEPES, 150 mM NaCl, 0.2 mM DTT, and 0.02% NP-40). The reaction was stopped with addition of 2X Laemmli sample buffer containing 10% β-mercaptoethanol.

Protein samples were heated at 95°C for 5 minutes before loading and separated onto SDS-PAGE cassette (Thermo Fisher Scientific Bolt™ 4-12% Bis-Tris Plus Gels) in MES running buffer. For direct protein detection, the SDS-PAGE gel was stained using InstantBlue® Protein Stain (Expedeon). For immunoblotting, the separated proteins were transferred onto polyvinylidene difluoride (PVDF) membrane using iBlot™ 2 Dry Blotting System (ThermoFisher Scientific). The blot was blocked with 5% skim milk in PBS containing 0.05% Tween-20 and probed with anti-mono or poly ADP-ribose binding reagents/antibodies MABE1076 (α-MAR), MABC547 (α-PAR), MABE1075 (α-MAR/PAR) (Millipore-Sigma) and anti-GST tag monoclonal antibody MA4-004 (ThermoFisher Scientific). The primary antibodies were detected with secondary infrared anti-rabbit and anti-mouse antibodies (LI-COR Biosciences). All immunoblots were visualized using Odyssey® CLx Imaging System (LI-COR Biosciences). The images were quantitated using Image J (National Institutes for Health (NIH)) or Image Studio software.

#### Kinetic analysis of ADP-ribose hydrolysis

To quantify the initial rate (*k*) of substrate decay associated with the four macrodomains, each data set represented in the substrate decay immunoblots in Fig. 6C, were fitted to a decaying exponential with the following functional form: ([S]_*initial*_-[S]_*final*_)*e*^(-[*k*/[S]_*initial*_)*t*]^+[S]_*final*_ (Mathematica 12, Wolfram Alpha). The decay plots and resulting values for the fitted parameter *k* along with statistic uncertainty (SD) are shown in Fig. 6D.

#### ELISA-based MAR hydrolysis

ELISA Well-Coated™ Glutathione plates (G-Biosciences, USA) were washed with phosphate-buffered saline (PBS) containing 0.05% Tween-20 (PBS-T) and incubated with 50 μL of 100 nM automodified MARylated PARP10-CD in PBS for one hour under room temperature. Following four washes with PBS-T, variable concentrations of SARS-CoV-2 Mac1 were incubated with MARylated PARP10-CD for 60 minutes at 37°C. Purified macrodomains were 2-fold serially diluted starting at 100 nM in reaction buffer prior to addition to MARylated PARP10-CD. Subsequently, ELISA wells were washed four times with PBS-T and incubated with 50 μL/well of anti-GST (Invitrogen MA4-004) or anti-MAR (Millipore-Sigma MAB1076) diluted 1:5,000 in 5 mg/ml BSA in PBS-T (BSA5-PBS-T) for 1 hour at room temperature. After four additional washes with PBS-T, each well was incubated with 50 μL diluted 1:5,000 in BSA5-PBS-T of anti-rabbit-HRP (SouthernBiotech, USA) or anti-mouse-HRP (Rockland Immunochemicals, USA) conjugate for 1 hour at room temperature. The plate was washed four times with PBS-T and 100 μL of TMB peroxidase substrate solution (SouthernBiotech, USA) was added to each well and incubated for 10 minutes. The peroxidase reaction was stopped with 50 μL per well of 1 M HCl before proceeding to reading. Absorbance was measured at 450 nm and subtracted from 620 nm using Biotek Powerwave XS plate reader (BioTek). As controls, MARylated PARP10-CD and non-MARylated PARP10 were detected with both anti-MAR and anti-GST antibodies. The absorbance of non-MARylated PARP10-CD detected with anti-MAR antibody was used to establish the background signal. The % signal remaining was calculated by dividing the experimental signal (+ enzyme) minus background by the control (no enzyme) minus the background.

### PAR Hydrolase Assay

#### Automodification of PARP1 protein

PARP1 was incubated with increasing concentrations of NAD^+^ to generate a range of PARP1 automodification levels. Highly purified human 6X-His-PARP1 (43) (5 μg) was incubated for 30 min at 30°C in a reaction buffer containing 100 mM Tris-HCl pH 8.0, 10 mM MgCl_2_, 10% (v/v) glycerol, 10 mM DTT, 0 to 500 μM NAD+, 10% (v/v) ethanol and 25 μg/mL calf thymus activated DNA (Sigma-Aldrich).

#### PARP1 ADP-ribose hydrolysis

To evaluate the PAR hydrolase activity of CoV macrodomains, 200 ng of slightly automodified PARP1 with 5 μM NAD^+^ or highly automodified with 500 μM NAD^+^ were used as substrates for the de-PARylation assays. Recombinant macrodomain protein (1 μg) was supplemented to the reaction buffer (100 mM Tris-HCl pH 8.0, 10% (v/v) glycerol and 10 mM DTT) containing automodified PARP1 and incubated for 1 hour at 37°C. Recombinant PARG (1 μg) was used as a positive control for PAR erasing (44). Reaction mixtures were resolved on 4–12% Criterion™ XT Bis-Tris protein gels, transferred onto nitrocellulose membrane and probed with the anti-PAR polyclonal antibody 96-10.

### Structure Determination

#### Crystallization and Data Collection

Purified SARS-CoV-2 Mac1 in 150 mM NaCl, 20 mM Tris pH 8.0 was concentrated to 13.8 mg/mL for crystallization screening. All crystallization experiments were setup using an NT8 drop-setting robot (Formulatrix Inc.) and UVXPO MRC (Molecular Dimensions) sitting drop vapor diffusion plates at 18°C. 100 nL of protein and 100 nL crystallization solution were dispensed and equilibrated against 50 μL of the latter. The SARS-CoV-2 Mac1 complex with ADP-ribose was prepared by adding the ligand, from a 100 mM stock in water, to the protein at a final concentration of 2 mM. Crystals that were obtained in 1-2 days from the Salt Rx HT screen (Hampton Research) condition E10 (1.8 M NaH_2_PO_4_/K_2_HPO_4_, pH 8.2). Refinement screening was conducted using the additive screen HT (Hampton Research) by supplementing 10% of each additive to the Salt Rx HT E10 condition in a new 96-well UVXPO crystallization plate. The crystals used for data collection were obtained from Salt Rx HT E10 supplemented with 0.1 M NDSB-256 from the additive screen (Fig. S1). Samples were transferred to a fresh drop composed of 80% crystallization solution and 20% (v/v) PEG 200 and stored in liquid nitrogen. X-ray diffraction data were collected at the Advanced Photon Source, IMCA-CAT beamline 17-ID using a Dectris Eiger 2X 9M pixel array detector.

#### Structure Solution and Refinement

Intensities were integrated using XDS (45, 46) via Autoproc (47) and the Laue class analysis and data scaling were performed with Aimless (48). Notably, a pseudo-translational symmetry peak was observed at (0, 0.31 0.5) that was 44.6% of the origin. Structure solution was conducted by molecular replacement with Phaser (49) using a previously determined structure of ADP-ribose bound SARS-CoV-2 Mac1 (PDB 6W02) as the search model. The top solution was obtained in the space group *P*21 with four molecules in the asymmetric unit. Structure refinement and manual model building were conducted with Phenix (50) and Coot (51) respectively. Disordered side chains were truncated to the point for which electron density could be observed. Structure validation was conducted with Molprobity (52) and figures were prepared using the CCP4MG package (53). Superposition of the macrodomain structures was conducted with GESAMT (54).

### Statistical Analysis

All statistical analyses were done using an unpaired two-tailed student’s t-test to assess differences in mean values between groups, and graphs are expressed as mean ±SD. Significant p values are denoted with *p≤0.05.

## ACCESSION CODES

The coordinates and structure factors for SARS-CoV-2 Mac1 were deposited to the Worldwide Protein Databank (wwPDB) with the accession code 6WOJ.

## ACKNOWLEDGEMENTS

We’d like to thank Ivan Ahel and Gytis Jankevicius (Oxford University) for providing protein expression plasmids; John Pascal (University of Montreal) and Marie-France Langelier (Universite de Montreal) for providing PARP1; and Wenqing Xu (University of Washington) for providing PARG. This research was funded by the National Institutes of Health (NIH) grant numbers P20 GM113117, P30GM110761, and AI134993-01, and University of Kansas start-up funds to A.R.F, and the Canadian Institutes of Health Research grant number MOP-418863 to G.G.P. Use of the IMCA-CAT beamline 17-ID at the Advanced Photon Source was supported by the companies of the Industrial Macromolecular Crystallography Association through a contract with Hauptman-Woodward Medical Research Institute. Use of the Advanced Photon Source was supported by the U.S. Department of Energy, Office of Science, Office of Basic Energy Sciences, under Contract No. DE-AC02-06CH11357.

## AUTHOR CONTRIBUTIONS

Conceptualization: ARF, YMOA, GGP

Data curation: YMOA, SL, JPG, ARF, EDH

Formal analysis: YMOA, DKJ, AR, SL, ARF, EDH

Funding acquisition: GGP, SL, ARF

Investigation: YMOA, MMK, AR, JPG, LN, PM, KPB

Methodology: YMOA, GGP, AR, JPG, EDH, SL, ARF

Project administration: GGP, SL, ARF

Resources: AR, SL, PG, ARF

Supervision: AR, GGP, SL ARF

Validation: YMOA, SL, AR, JPG, GGP, ARF

Visualization: YMOA, ARF, AR, SL, JPG

Writing – original draft: YMOA, SL, ARF

Writing – review & editing: all authors

## SUPPLEMENTAL FIGURE LEGENDS

**Figure S1**. Purification and crystallization of macrodomain proteins. **A)** Macrodomain proteins were purified as described in Methods. Equimolar amounts of the recombinant proteins were run on a polyacrylamide gel and visualized by Coomassie staining. **B)** Crystals of SARS-CoV-2 Mac1 obtained with Salt Rx HT E10 supplemented with 0.1 M NDSB-256.

**Figure S2.** Extended residues at the C-terminus of the SARS-CoV-2 Mac1 clashed with symmetry related molecules. **A)** Comparison of the amino acid sequence of SARS-CoV-2 Mac1, 6W02 and 6WEY. **B)** Superposition of SARS-CoV-2 Mac1 (magenta) subunit B onto subunit A of 6W02 reveals that the C-terminus would clash with symmetry related molecules (coral).

**Figure S3**. Comparison of the SARS-CoV-2 Mac1 protein with homologous structures. **A-B)** Superposition of SARS-CoV-2 Mac1 (magenta) with other recently determined homologous structures. **A)** SARS-CoV-2 Mac1 apo structure (6WEN), **B)** SARS-CoV-2 Mac1 complexed with ADP-ribose (6W02). The ADP-ribose molecule is colored gray for SARS-CoV-2 and is represented as green cylinders for 6W02 in panel **B**. **C-D)** Comparison of the residues in the ADP-ribose binding site. **C**) SARS-CoV-2 Mac1 apo structure (blue, 6WEN), **D**) SARS-CoV-2 Mac1 complexed with ADP-ribose (green, 6W02). The ADP-ribose of SARS-CoV-2 is rendered as gray cylinders, and is represented as green cylinders for 6W02 in panel **B**.

**Figure S4**. ADP-ribose binding of macrodomain proteins by DSF assay. The macrodomain proteins (10 μM) were incubated with increasing concentrations of ADP-ribose and measured by DSF as described in Methods. Mdo2 n=4; SARS-CoV n=6; MERS-CoV n=5; SARS-CoV-2 n=3.

**Figure S5**. Affinity of ADP-ribose binding antibodies for ADP-ribosylated PARP10 CD. MARylated PARP10 and non-MARylated PARP10 CD were detected by immunoblot (IB) with anti-GST (Invitrogen, MA4-004), anti-ADP-ribose binding reagents: anti-MAR (Millipore-Sigma MAB1076), anti-PAR (Millipore-Sigma MABC547), and anti-MAR/PAR (Millipore-Sigma MABE1075) antibodies.

**Figure S6.** MARylated PARP10 stability over time. The presence of mono-ADP-ribose of automodified PARP10 without any macrodomain was detected at 6 time points by immunoblot analysis with the anti-GST (Invitrogen, MA4-004) and anti-ADP-ribose binding reagent anti-MAR (Millipore-Sigma MAB1076).

**Figure S7.** The CoVs and human Mdo2 macrodomain proteins were incubated with MARylated PARP10 CD *in vitro* for the indicated times at 37°C. Total PARP10 CD and macrodomain protein levels were determined by Coomassie Blue (CB) staining. Results showone experiment of three independent experiments.

**Figure S8**. Differential PARylation of PARP1 by varying concentrations of NAD^+^. Recombinant human PARP1 was automodified in a reaction buffer supplemented with increasing concentrations of NAD^+^ to generate substrates for the PAR hydrolase assays. PAR was detected by immunoblot analysis of reaction products with the anti-PAR antibody 96-10.

