## Supplemental Figures for "The SARS-CoV-2 conserved macrodomain is a mono-ADP-ribosylhydrolase"

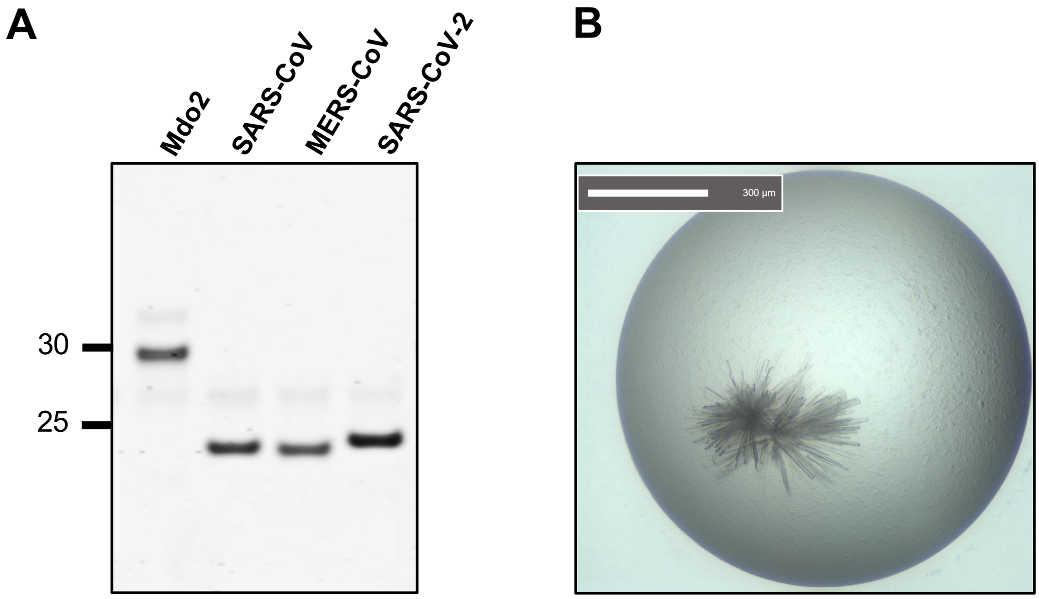
A
B
Mdo2
SARS-CoV
MERS-CoV
SARS-CoV-2
30
25
300 µm

**A**

```
                   0         10        20        30        40
                   |....|....|....|....|....|....|....|....|....|....
SARS-CoV-2 Mac1    GIEVNSFSGYLKLTDNVYIKNADIVEEAKKVKPTVVVNAANVYLKHGGGV
6W02               -GEVNSFSGYLKLTDNVYIKNADIVEEAKKVKPTVVVNAANVYLKHGGGV
6WEY               --GVNSFSGYLKLTDNVYIKNADIVEEAKKVKPTVVVNAANVYLKHGGGV

                   50        60        70        80        90
                   |....|....|....|....|....|....|....|....|....|....
SARS-CoV-2 Mac1    AGALNKATNNAMQVESDDYIATNGPLKVGGSCVLSGHNLAKHCLHVVGPN
6W02               AGALNKATNNAMQVESDDYIATNGPLKVGGSCVLSGHNLAKHCLHVVGPN
6WEY               AGALNKATNNAMQVESDDYIATNGPLKVGGSCVLSGHNLAKHCLHVVGPN

                   100       110       120       130       140
                   |....|....|....|....|....|....|....|....|....|....
SARS-CoV-2 Mac1    VNKGEDIQLLKSAYENFNQHEVLLAPLLSAGIFGADPIHSLRVCVDTVRT
6W02               VNKGEDIQLLKSAYENFNQHEVLLAPLLSAGIFGADPIHSLRVCVDTVRT
6WEY               VNKGEDIQLLKSAYENFNQHEVLLAPLLSAGIFGADPIHSLRVCVDTVRT

                   150       160       170
                   |....|....|....|....|....|
SARS-CoV-2 Mac1    NVYLAVFDKNLYDKLVSSFLEMKSEK
6W02               NVYLAVFDKNLYDKLVSSFLE-----
6WEY               NVYLAVFDKNLYDKLVSSFLEMKS--
```

**B**

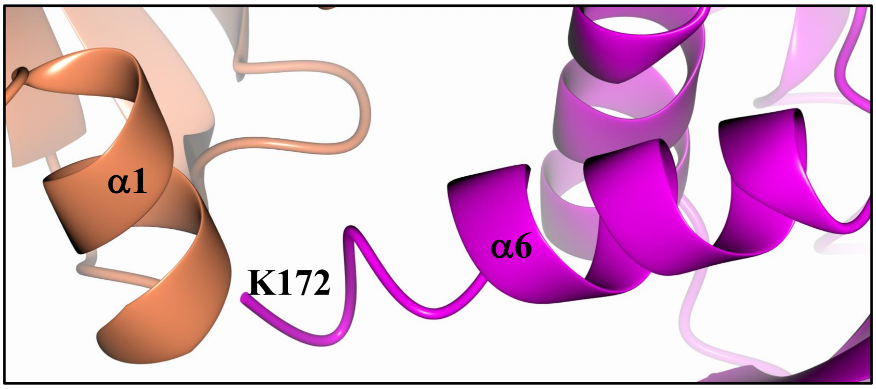

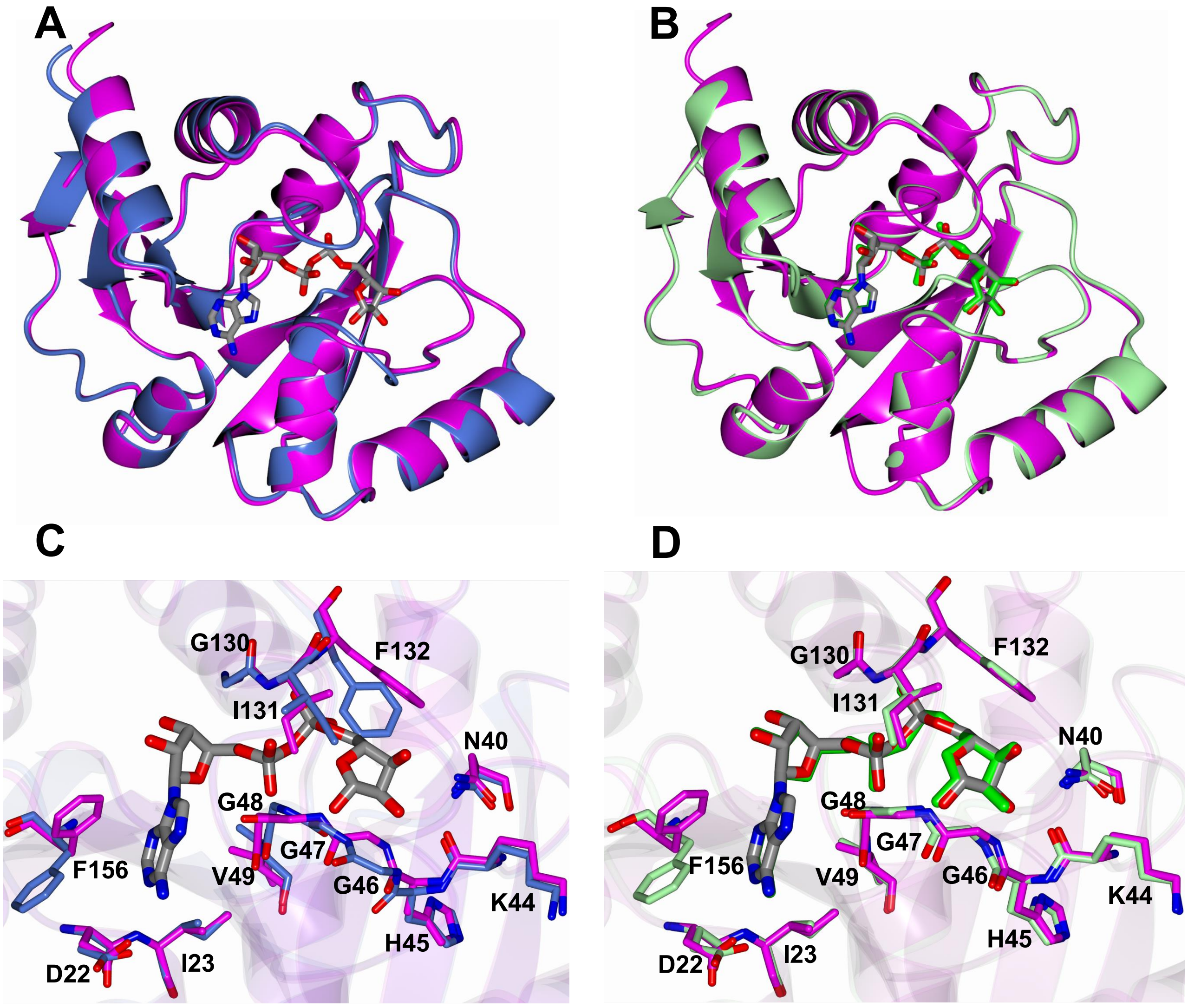

A
B
C
D
G130
F132
I131
N40
G48
G47
F156
V49
G46
K44
H45
D22
I23
G130
F132
I131
N40
G48
G47
F156
V49
G46
K44
H45
D22
I23

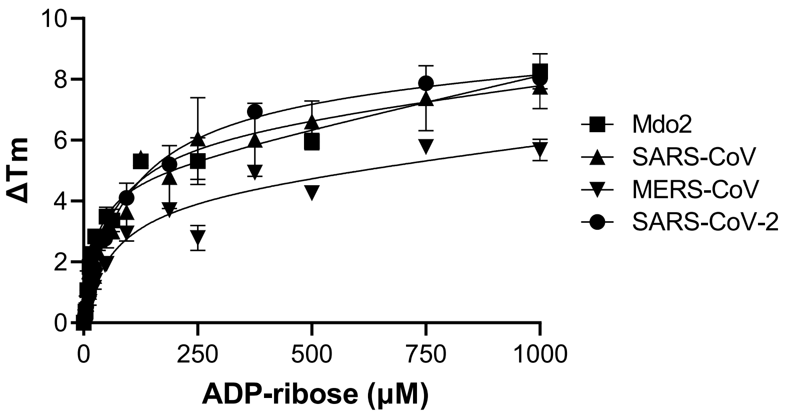

10
8
6
4
2
0
ΔTm
0
250
500
750
1000
ADP-ribose (μM)
Mdo2
SARS-CoV
MERS-CoV
SARS-CoV-2

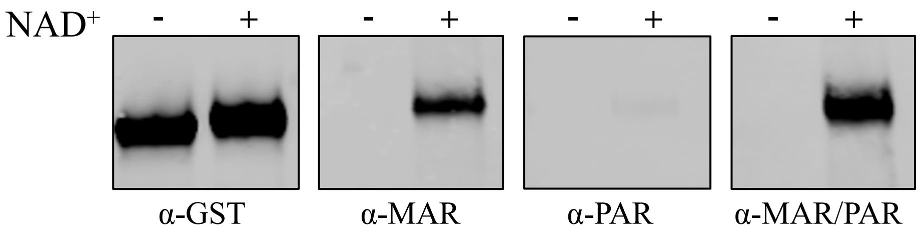

NAD+
-
+
-
+
-
+
-
+
α-GST
α-MAR
α-PAR
α-MAR/PAR

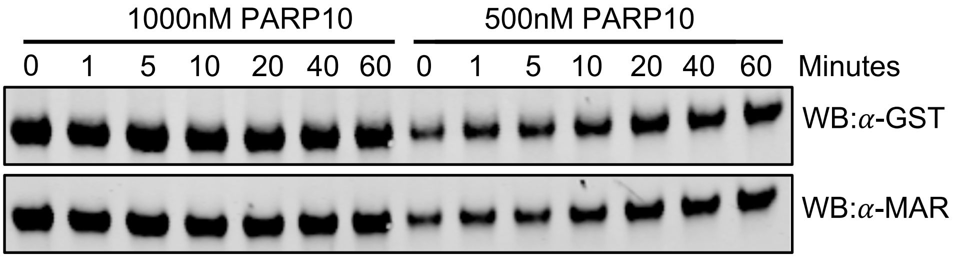

1000nM PARP10
500nM PARP10
0 1 5 10 20 40 60 0 1 5 10 20 40 60 Minutes
WB:α-GST
WB:α-MAR

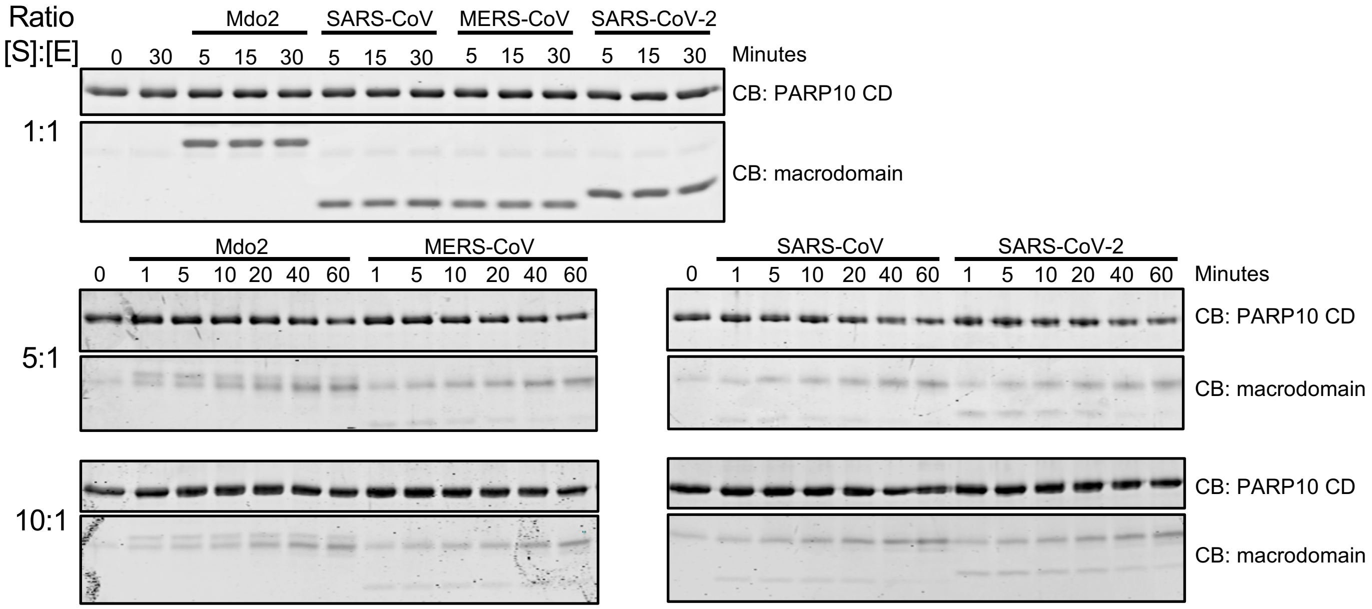

Ratio
[S]:[E]
Mdo2
SARS-CoV
MERS-CoV
SARS-CoV-2
0 30 5 15 30 5 15 30 5 15 30 5 15 30
Minutes
CB: PARP10 CD
1:1
CB: macrodomain
Mdo2
MERS-CoV
0 1 5 10 20 40 60 1 5 10 20 40 60
SARS-CoV
SARS-CoV-2
0 1 5 10 20 40 60 1 5 10 20 40 60
Minutes
CB: PARP10 CD
5:1
CB: macrodomain
CB: PARP10 CD
10:1
CB: macrodomain

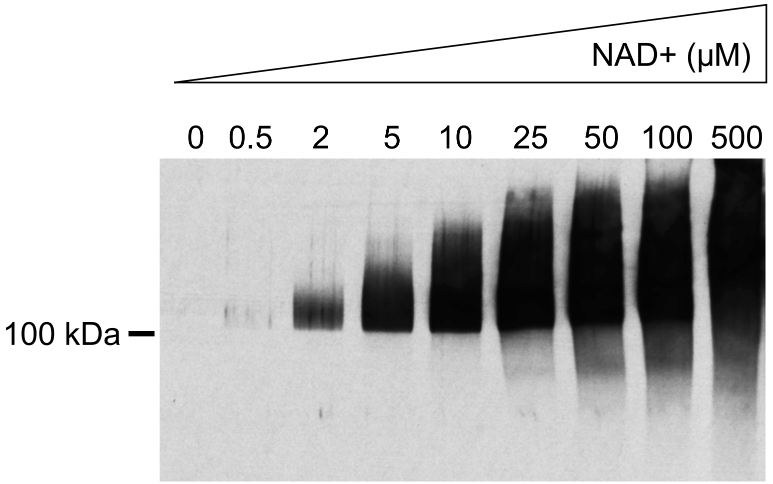
NAD+ (µM)
0
0.5
2
5
10
25
50
100
500
100 kDa
